# Pro-inflammatory microglia drive escalated alcohol consumption during early abstinence

**DOI:** 10.64898/2026.07.30.741570

**Authors:** PE Anton, BM Materia, DF Lovelock, SA McDonald, SP Lyons, A Mordant, LE Herring, J Besheer, LG Coleman

## Abstract

Despite growing evidence that neuroimmune dysfunction contributes to Alcohol Use Disorder (AUD) pathology, the underlying neuroinflammatory mechanisms that may promote alcohol consumption are not as clear. We recently report that specific knockdown of interferon regulatory factor 7 (IRF7) in the anterior insula (aIC) mitigates escalation in ethanol self-administration in rats. In addition, we find pro-inflammatory activation of microglia contributes to other AUD-related behavioral impairments. Here, we sought to determine if pro-inflammatory activation of microglia from ethanol contributes to elevations in IRF7 and ethanol self-administration in rats. Male Wistar rats were trained under our ethanol self-administration paradigm (15% v/v; FR2 vs inactive lever) followed by 1-4 cycles of chronic intermittent ethanol vapor exposure (CIE). To inhibit microglia, rats were treated with minocycline (30mg/kg, i.p.) before and after each ethanol vapor session. Escalation in self-administration and biochemical markers were assessed 72 hours into abstinence. We found CIE increased ethanol self-administration, which was positively correlated with aIC IRF7 levels. Minocycline treatment blunted IRF7 expression and alleviated ethanol self-administration following CIE. LC-MS/MS proteomics of primary microglia from the aIC of rats following CIE with and without minocycline treatment indicate minocycline promotes metabolic and ribosomal re-wiring of microglia during early abstinence. These data suggest a role for microglia in driving both IRF7 levels and escalation in ethanol self-administration in early abstinence.

## Introduction

Alcohol Use Disorder (AUD) currently effects nearly 28 million people in the United States^1^. Past heavy alcohol use is a significant risk factor for AUD^2^, which is characterized by cycles of heavy alcohol exposure, short-term withdrawal from alcohol, and a return to drinking^3^. The withdrawal phase (i.e. early abstinence) includes negative affect and alcohol cravings, and the severity of these symptoms can predict a return to use^4^. Therefore, early abstinence represents a key phase for therapeutic intervention. However, the underlying cellular and molecular mechanisms by which previous heavy alcohol exposure followed by early abstinence promotes AUD are complex. Heightened neuroimmune activation is a contributing factor to AUD-related neuronal dysfunction and behavioral impairments^5–7^. Human post-mortem brain tissue from subjects with AUD express increased innate immune factors including glial activation markers^8^, Toll like receptors (TLRs)^9^, and danger associated molecular patterns (DAMPs)^10^. These features have been replicated using rodent models of binge or chronic ethanol exposure^11,12^. We, and others, show that microglia, resident brain macrophages, play a vital role in orchestrating neuroinflammation from ethanol exposure and associated AUD-related behaviors in abstinence^11,13^. For example, we report that repeated binge ethanol exposure promotes cortical pro-inflammatory cytokine induction and ethanol-induced negative affect in mice several weeks into abstinence, which is mitigated by inhibition of pro-inflammatory microglia ^11^. Moreover, pharmacological depletion of microglia reduces escalation of ethanol intake in mice^14^ and minocycline, a drug that prevents the pro-inflammatory activation of microglia^15^, reduces ethanol intake in the 2-bottle choice task in mice^16^.

These studies implicate microglia in ethanol consumption and suggest microglia help initiate innate immune cascades in AUD. However, the specific innate immune factors that microglia induce to promote ethanol-intake are not as well defined. We previously find that chronic intermittent ethanol vapor exposure (CIE) induces interferon regulatory factor 7 (IRF7), a master regulator of pro-inflammatory interferon signaling^13,17^. IRF7 is a pro-inflammatory transcription factor that is primarily in neurons and is downstream of several TLRs, such as Toll-like receptor-7 (TLR7). Specific knock down of IRF7 in the anterior insula (aIC), a pertinent brain region relevant to ethanol consumption, prevents escalation of ethanol self-administration following CIE ^13^. These data suggest that targeting neuroimmune activation can alleviate AUD-related behaviors. Further, we reported previously that microglia release pro-inflammatory factors capable of inducing neuronal TLR7^18,19^. However, the interaction between microglia activation, IRF7 signaling, and ethanol consumption is not understood. Therefore, we investigated if inhibiting pro-inflammatory microglia can prevent ethanol-induced expression of IRF7 and escalation of ethanol-self administration. We then performed LC-MS/MS proteomics on primary aIC microglia from rats in early abstinence following ethanol and/or minocycline treatment to reveal the neuroprotective mechanisms of microglia during early abstinence.

## Results

### Minocycline Reduces IRF7 and Ethanol-Self Administration Following CIE

The risk of relapse is particularly high during the first 90 days of abstinence^4^. In rodent models of AUD, ethanol self-administration is often measured up to 24 hours following previous ethanol exposure^20,21^, during acute detoxification. Therefore, the underlying molecular factors that may encourage ethanol consumption past the acute detoxification phase and into early abstinence are not well understood. We recently reported that chronic intermittent ethanol (CIE) exposure increases ethanol self-administration in female rats beyond the initial detoxification period at 72 hours into abstinence^13^. Here we explore ethanol self-administration in early abstinence in males. Male Wistar rats underwent four cycles CIE with a week of operant self-administration interleaved between CIE cycles (Figure 1A). Similar to our findings in females, CIE caused a robust enhancement of operant self-administration of alcohol in male rats 72h into early abstinence (Figure 1B).

**Figure 1.**
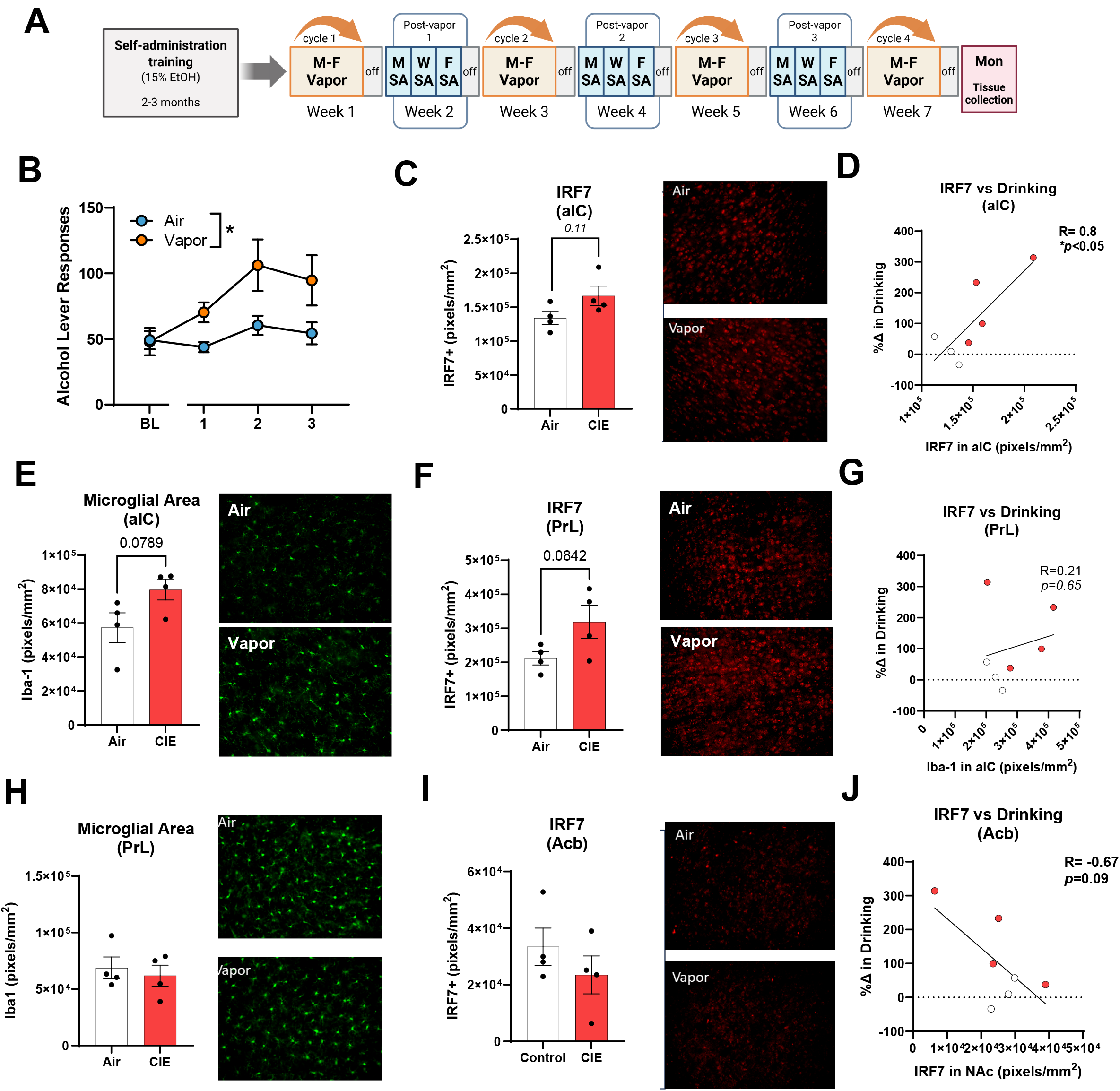
IRF7 is positively correlated with ethanol self-administration in the anterior insula. (A) Male Wistar rats underwent our ethanol self-administration training program followed by 4 cycles of CIE and ethanol self-administration. Tissue was collected 72 hours into abstinence. (B) Average number of ethanol lever responses during each ethanol self-administration week. (C) IRF7 expression in the aIC following our CIE and ethanol self-administration regime. (D) Correlations between changes in ethanol self-administration compared to baseline and aIC IRF7 levels. (E) Iba-1 area in the aIC (F) IRF7 area in the PrL (G) correlation between IRF7 expression and change in ethanol consumption in the PrL (H) Iba-1 staining area in the PrL (I) IRF7 expression in the Acb (J) correlation between IRF7 and ethanol consumption in the Acb. Red dots indicate CIE, white dots indicate Air group. p values are indicated by student’s t-test or spearman’s correlation. N=3-4 per group.

Our prior report found that dependence-induced escalation of drinking involves induction of neuronal IRF7 in the anterior insula (aIC) of female mice. We also previously report that pro-inflammatory microglia are necessary for downstream neuroimmune dysfunction from ethanol exposure^11,22^. Therefore, we assessed IRF7 and microglial morphology in the aIC and two additional regions associated with AUD, the prelimbic cortex (PrL) and the nucleus accumbens (Acb)^3^. CIE caused a slight increase in IRF7 in the aIC that approached statistical significance (Figure 1C). However, a strong positive correlation was found between aIC IRF7 and alcohol self-administration (Figure 1D, R=0.8, \**p<*0.05). A trend toward an increase in microglial cell area was also found in the aIC (Figure 1E, *p=*0.079) with no changes in microglial number (Supplemental Figure 1A). Similar to IRF7, a trend toward a positive correlation between aIC microglial area and alcohol intake was found (Supplemental Figure 1B, R=0.61, *p=*0.15). Similar to our report in females^13^, a trend toward an increase in IRF7 was also seen in the PrL (Figure 1F, *p=*0.08). However, as we reported in females, IRF7 in the PrL was not correlated with drinking (Figure 1G), and there was no change in microglial area (Figure 1H). In the Acb IRF7 was not changed and a trend toward an inverse correlation between IRF7 and ethanol intake was found (Figure 1J). Thus, increases in IRF7 and microglia area in the aIC were positively associated with escalation of alcohol intake after CIE, consistent with findings in females^13^.

### Minocycline Reduces IRF7 in the aIC and Acb after CIE

Pro-inflammatory microglia are key initiators of ethanol-induced neuroimmune dysfunction^11,22^. Therefore, we next assessed the impact of blocking pro-inflammatory microglial polarization on CIE-induced increases in IRF7 expression, which is primarily neuronal. Rats were given minocycline, an anti-inflammatory drug that blunts microglia activation^15^. Minocycline (30mg/kg, i.p.) or vehicle were given immediately prior to and after each ethanol vapor session (Figure 2A). IRF7 and microglial area were assessed 72 hours after final ethanol vapor exposure, the same timepoint where we observed CIE-related inductions in ethanol self-administration (Figure 1B). With this single week of CIE, we did not find changes in microglial area in the aIC or Acb (Figure 2B-C). However, increases in IRF7 were found. In the aIC minocycline caused a significant decrease in IRF7 compared to CIE alone (Figure 2D, \*\**p*<0.01). With this single week of CIE, an increase in IRF7 was found in the Acb CIE (Figure 2E, p<0.05), versus the lack of a change seen after 4 cycles, suggesting an adaptative response with multiple CIE cycles. Similar to the aIC, minocycline blocked this increase in IRF7 in the Acb (Figure 2C). Thus, inhibition of pro-inflammatory microglial activation blunts IRF7 induction in the aIC and Acb.

**Figure 2.**
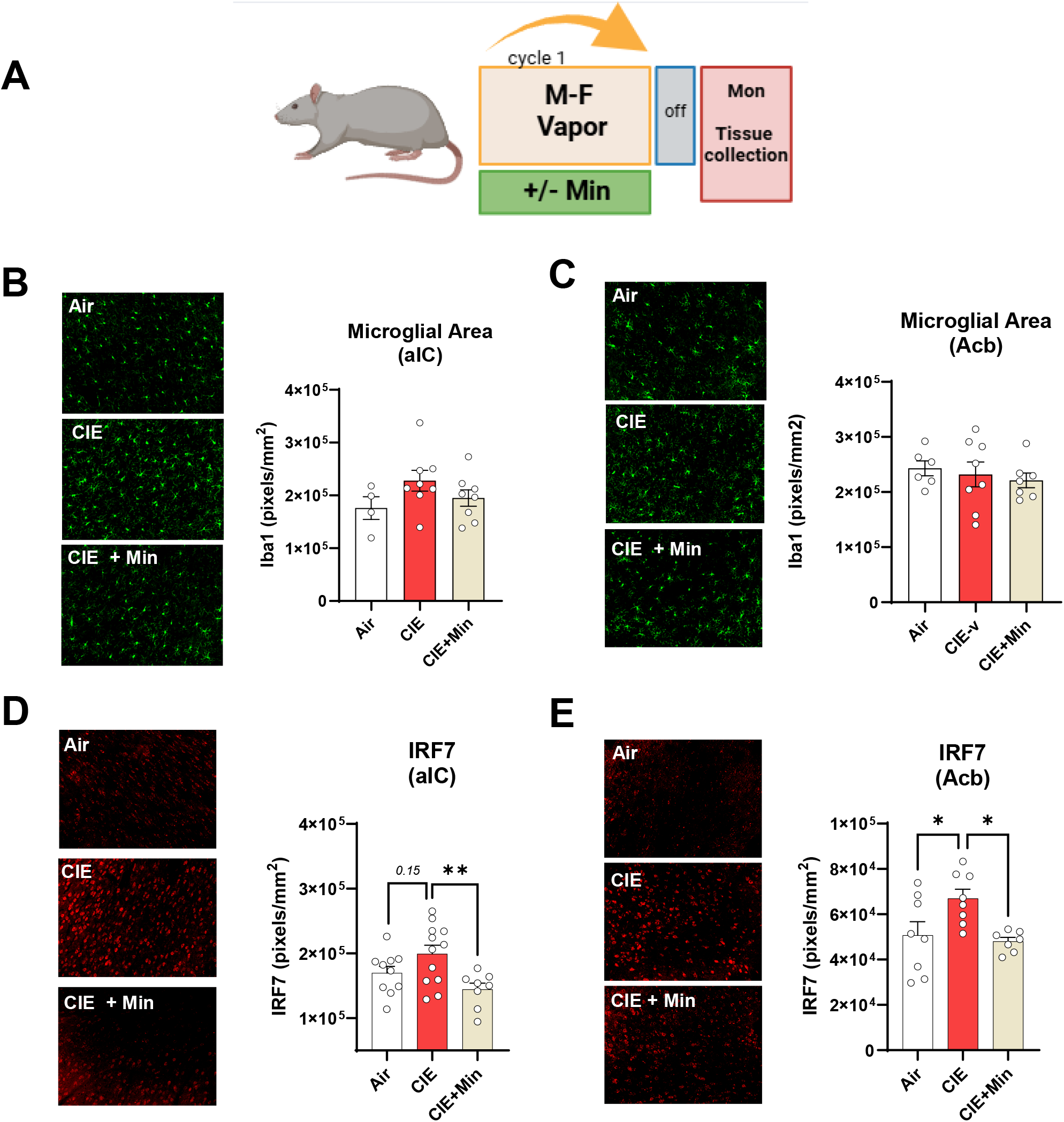
Minocycline reduces CIE-related IRF7 expression. (A) Male Wistar rats were exposed to one cycle of CIE. Minocycline (30 mg/kg, i.p.) or a vehicle were administered before and after each vapor session. Tissue was collected 72 hours after final vapor session. Immunofluorescence of Iba-1 in (B) the aIC and (C) the Acb. Immunofluorescence of IRF7 in (D) the aIC and (E) the Acb. N=4-8 per group, *\*p<0*.*05, **p<0*.*01* by one-way ANOVA.

### Minocycline prevents ethanol-induced microglial cellular stress phenotypes

Although we did not observe microglial morphological changes due to one cycle of CIE with or without minocycline (Figure 2B-C), we did find a minocycline-related effect on IRF7. These data suggest there may be underlying molecular effects of CIE on microglia that are prevented by minocycline. To investigate these molecular changes, we characterized aIC microglia from rats in the early abstinence phase following CIE and or minocycline treatment using LC-MS/MS proteomics (Figure 3A). The aIC was chosen due to the significant correlations between aIC IRF7 and ethanol escalation (Figure 1D). We observed an enrichment in microglia-specific proteins following CD11b^+^ selection and LC-MS/MS (Supplemental Figure 2A). We observed clustering between our biological replicates, although did not see separation between groups using principal component analysis (Supplemental Figure 2B). In microglia from ethanol exposed rats compared to control (EtOH vs. Air) 26 proteins were significantly altered and in primary microglia from rats receiving minocycline and ethanol compared to microglia from ethanol exposed rats (CIE+Min vs. CIE), we identified 33 differently expressed proteins (DEPs) (p<0.05, log_2_FC abs >0.5) (Figure 3B-C). Using Qiagen Ingenuity Pathway Analysis (IPA) we identified 199 canonical pathways that were significantly altered in microglia from ethanol exposed rats compared to control (p≤0.05, z score abs≥2). We specifically find predicted increases in pro-inflammatory canonical pathways due to CIE as evidenced by increased NLR signaling, Class I MHC antigen presentation, and TLR and IL-1 signaling. The induction of these pathways by CIE were inhibited by minocycline (Figure 3D). These data suggest that while one cycle of CIE is not sufficient to induce morphological features of microglia activation, it does promote innate immune signaling in microglia, which is prevented by minocycline.

**Figure 3.**
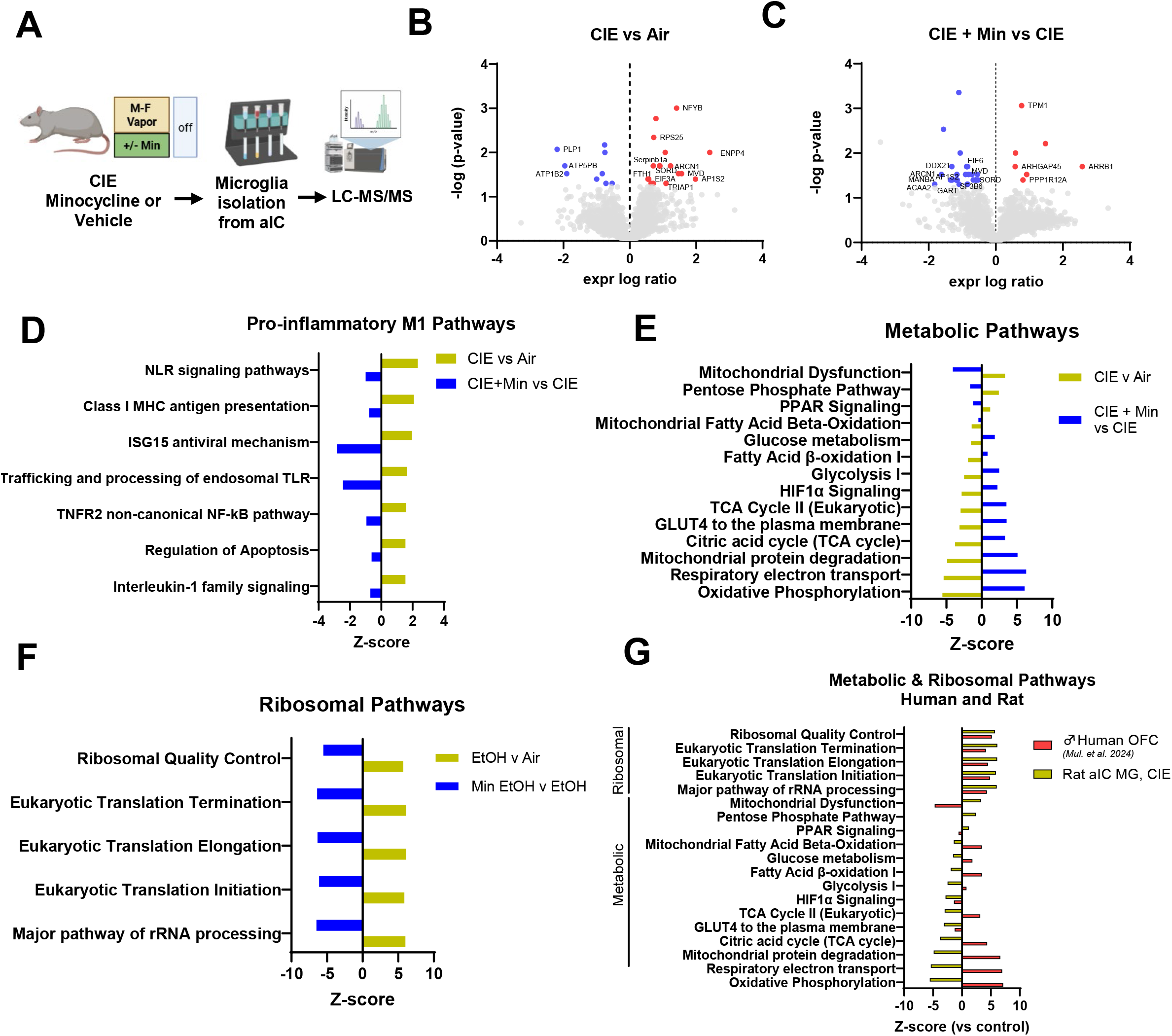
Minocycline normalizes ethanol-Induced microglial stress phenotypes. (A) Rats were exposed to one cycle of CIE minocycline treatment or vehicle. 72 hours into abstinence microglia were isolated from the aIC for LC-MS/MS proteomics. Differentially expressed proteins in primary microglia from (B) CIE vs Air and (C) CIE+Min vs CIE comparisons. Blue dots indicate significantly reduced proteins (p<0.05, log_2_fc <-0.5) and red represents significantly increased proteins (p<0.05, log_2_fc>0.5). (D) Pro-inflammatory IPA canonical pathways upregulated by CIE (gold bars) and reduced by the addition of minocycline (blue bars). Additional IPA pathways showing (E) metabolic and (F) ribosomal pathways in the CIE and CIE + Min groups. (G) Comparison of ribosomal and metabolic pathways from aIC microglia from CIE rats and bulk proteomics tissue from human AUD brain (Mulholland et al. 2024). N=4 per group. All canonical pathways shown are *p<0*.*05*.

We then investigated other microglia phenotypes induced by CIE or minocycline to find potential regulators of microglial inflammatory signaling. Further exploration of ethanol-related DEPs suggest a microglial phenotype associated with increased protein synthesis and processing (EIF3A, RPS25, SNRPA, ARCN1, AP1S2), cellular stress responses (FTH1, TRIAP1, SERPINB1A) and a shift away from oxidative phosphorylation (MVD, SORD, ATP5B) (Figure 3B). In primary microglia from rats receiving minocycline and ethanol compared to microglia from ethanol exposed rats (CIE+Min vs. CIE) DEPs were associated with increased cellular motility and migration (ARRB1, ARHGAP45, PPP1R12A, TPM1), reduced protein synthesis and processing (EIF6, ARCN1, AP1S2, DDX21, SF3B6) and reduced lysosome machinery (MANBA, GALC, M6PR, CTSK, PLBD2) (Figure 3C). Given these changes, we further interrogated the influence of minocycline and/or ethanol on the metabolic and ribosomal state of microglia. We find that canonical pathways associated with mitochondria dysfunction are increased due ethanol exposure, while other energetic pathways including the TCA cycle, glycolysis, and oxidative phosphorylation are decreased. Minocycline prevents these ethanol-related deficits (Figure 3E). Despite the predicted suppression of cell metabolism due to ethanol, ribosomal and translation pathways were predicted to be increased by CIE with reversal by minocycline (Figure 3F). These data suggest abstinence from ethanol leaves microglia in a metabolically stunted state but translationally active state, associated with pro-inflammatory signaling. These data further suggest minocycline prevents CIE-induced microglial dysfunction through metabolic and/or translational re-programming. We then compared our data to human, using the proteome of the orbitofrontal cortex (OFC) from AUD postmortem tissue from male donors^23^. Several ribosomal pathways, particularly involving translation, were positively regulated in bulk OFC tissue due to AUD in a similar manner to CIE (Figure 3G). However, bulk tissue from AUD subjects displayed different metabolic pathway profiles than primary microglia from ethanol exposed rats (Figure 3G).

Consistent with the DEPs, the top 5 suppressed IPA canonical pathways due to ethanol exposure were metabolic (i.e., oxidative phosphorylation, respiratory electron transport chain, mitochondrial protein degradation) and the most strongly predicted upregulated pathways due to ethanol involved translation (processing of mRNA, translation initiation and elongation, rRNA processing) (Figure 4A). We find the opposite pattern when performing the CIE+Min vs. CIE comparison (Figure 4B). Oxidative phosphorylation pathways were amongst the top upregulated pathways, whereas translation-related pathways were amongst the most strongly predicted down regulated canonical pathways (Figure 4B). When comparing microglia from CIE+Min vs. Air, these features remain consistent (Supplemental Figure 2E-F) and distinct from the Min vs. Air comparison (Supplemental Figure 2G-H). Gene ontology (GO) analysis also indicates a CIE-related increase in translation and suppressed oxidative phosphorylation, in addition to suppressed pathways relating to synaptic vesicle regulation (Figure 4C). GO analysis of CIE + Min vs CIE comparison suggest suppressed translation, inflammatory responses, and glycolipid metabolism. CIE + Min was predicted to increase synapse regulation, oxidative phosphorylation, and cell motility compared to CIE (Figure 4D), consistent with the DEPs in Figure 3B.

**Figure 4.**
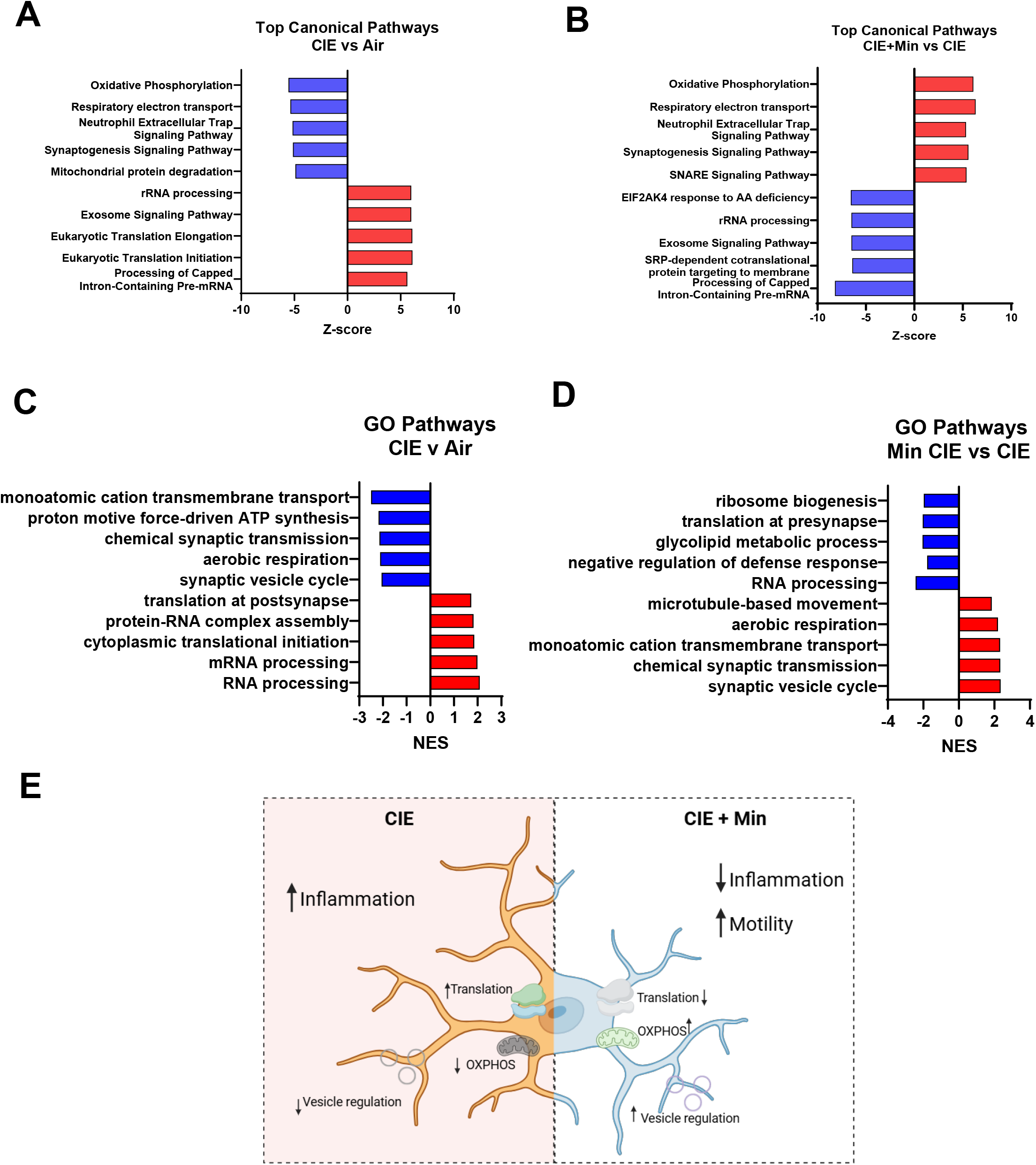
Minocycline predicted to promote microglia remodeling functions after CIE. Top 5 increased (red bars) and top 5 decreased (blue bars) IPA canonical pathways from the (A) CIE vs. Air and (B) CIE + Min vs CIE comparisons. GO analysis of the top 5 increased (red bars) and top 5 decreased (blue bars) pathways due to (C) CIE or (D) CIE + Min. N=4 per group. (E) Summary of findings from proteomics. All pathways shown are *p<0*.*05*.

On a molecular level, these data suggest microglia from CIE rats exhibit metabolic dysfunction and proteostasis distress, a phenotype consistent with microglial inflammation^24,25^. Moreover, GO analysis suggests these underlying features may result in impaired synaptic regulation in the aIC. However, the proteome of microglia from CIE + Min rats suggests microglia are more metabolically active and less inflammatory. Functionally, GO analysis suggests minocycline during CIE promotes a microglial movement and synaptic regulation (Figure 4E).

### Minocycline prevents CIE-induced escalation in alcohol self-administration

Given that LC-MS/MS suggests improved microglial function by minocycline, and because minocycline reduced aIC IRF7 levels, we next assessed the impact of minocycline on CIE-induced escalation of drinking. Rats were trained in operant self-administration of ethanol followed by one week of CIE with or without minocycline (Figure 5A, Supplemental Figure 3A). One week of CIE caused a robust increase in alcohol-self administration, increasing lever presses by 30% at 72h and 5 days into abstinence (Figure 5B). Minocycline treatment abolished the CIE-induced escalation in self-administration, with alcohol lever presses remaining near pre-CIE baseline levels (Figure 5B-C). Minocycline did not impact locomotion or total body weight indicating this suppression was not due to sedation or malaise (Figure 5D, Supplemental Figure 3B). Reductions in ethanol-self administration were associated with reduced microglia area in the aIC and Acb with minocycline (Figure 5E-F). Strong positive correlations were found between microglial area and drinking escalation in the aIC (R=0.73, \*\**p<*0.01) and Acb (R=0.57, \**p*<0.05) (Figure 3G-H). Interestingly, in this cohort no change in IRF7 was seen in the aIC with minocycline (Figure 5F), though a significant reduction was found in the Acb (Figure 5J). However, there were significant positive correlation between IRF7 with ethanol intake in the aIC (Figure 5K, R=0.67, *p<0*.*05*) with a trend in the Acb (R=0.4, *p=*0.14, Figure 5L), We did not observe differences in FosB, a marker of neuronal activation in the aIC or Acb due to minocycline treatment (Supplemental Figure 3C-D), nor differences in the neurotrophic factor BDNF (Supplemental Figure 3E-F). These data suggest preventing microglial pro-inflammatory signaling during CIE is a strategy to prevent escalation in ethanol-self administration.

**Figure 5.**
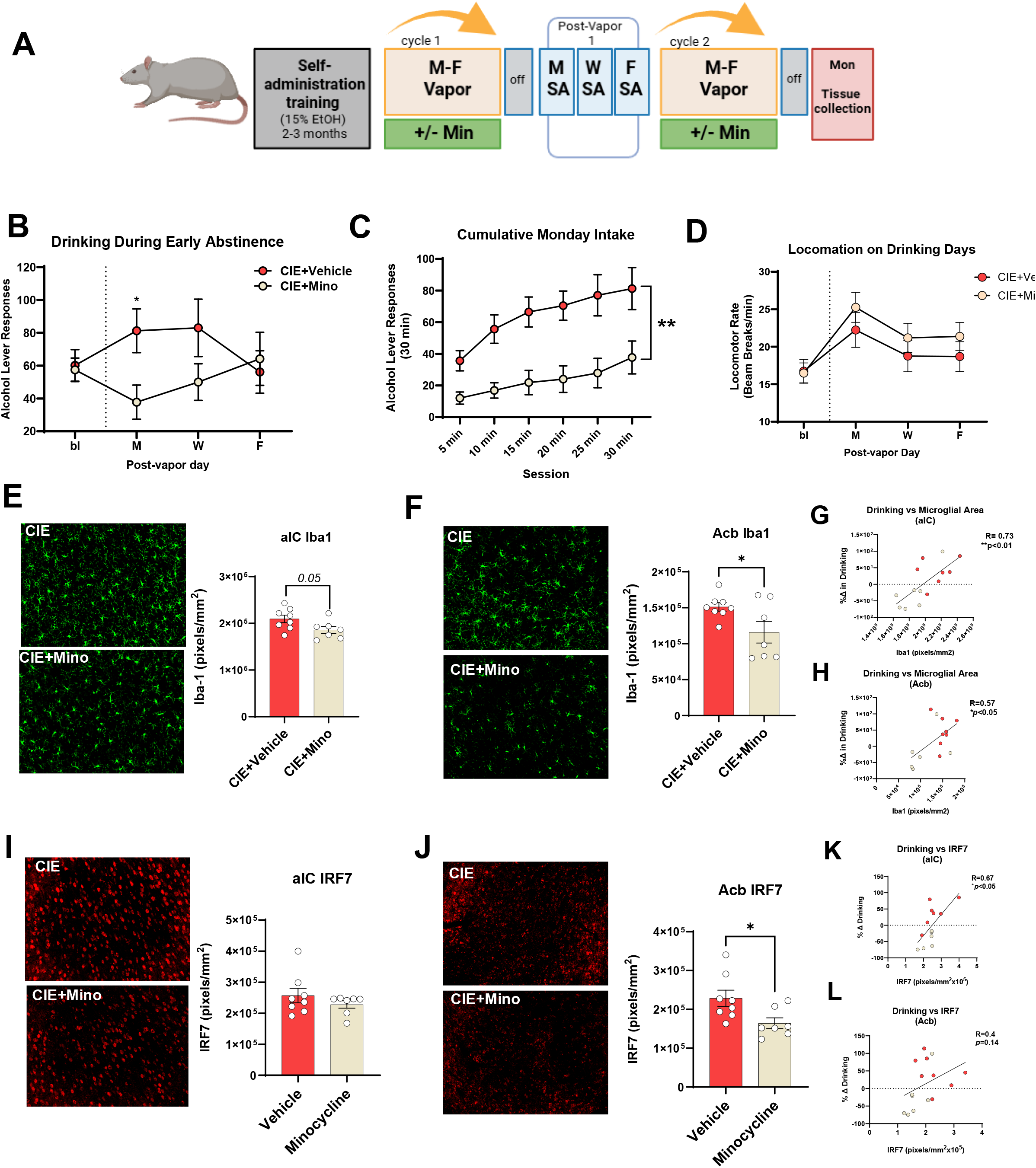
Minocycline Reduces Ethanol Self-Administration Following Chronic Intermittent Ethanol Exposure. (A) Rats underwent ethanol self-administration training followed by CIE +/-minocycline treatment and ethanol self-administration sessions. (B) Average number of ethanol lever responses during each ethanol self-administration session. (C) cumulative ethanol lever presses during the first 30minutes of the initial ethanol-self administration session. (D) Total number of beam breaks per minute during each ethanol self-administration session. (E) Iba-1 expression and (F) IRF7 expression in the aIC following CIE +/-minocycline. Correlations between aIC Iba-1 (G) and IRF7 (H) levels and ethanol consumption. (I) Iba-1 and (J) IRF7 expression in the NAc and correlations between NAc Iba-1 (K) and IRF7 (L) levels and ethanol consumption. Red dots indicate CIE, gold dots indicate CIE + Min. N=4/group. \**p<0*.*05* by student’s t-test.

## Discussion

Early abstinence from alcohol is a critical window for intervention to prevent continued drinking^4,26^. Therefore, understanding the underlying molecular neurobiology of early abstinence that promotes alcohol consumption may reveal novel translational targets for AUD. From rodent models of binge or heavy alcohol exposure, we and others report pro-inflammatory activation of microglia and downstream innate immune signaling drives neuronal injury during acute ethanol detoxification (i.e. up to 24 hours following final ethanol exposure)^18,27^. More recently, we reported pro-inflammatory microglia mediate long-lasting negative affect behaviors from binge ethanol exposure (i.e., up to nearly 6 weeks in abstinence), suggesting microglia are key mediators of AUD-related behaviors^11^. However, microglial phenotypes and their effect on ethanol consumption during early abstinence are not as well described. Here, we show that CIE promotes ethanol self-administration during early abstinence in male rats with correlative increases in aIC IRF7 expression. Intervention with minocycline, a drug that inhibits pro-inflammatory activation of microglia, reduces both IRF7 expression and ethanol self-administration due to CIE, suggesting microglia promote drinking behaviors in early abstinence. LC-MS/MS profiling of the microglial proteome during early abstinence suggests CIE induces an inflammatory phenotype of microglia, associated with a proteome signature of suppressed cell metabolism and a proteostasis stress. Minocycline treatment at the time of CIE prevented these features and promoted markers of cell motility and synapse regulation.

We found that ethanol self-administration was positively correlated with IRF7 levels in the aIC. These data are consistent to our previous findings that CIE-induced IRF7 in the aIC promotes ethanol consumption in female rats^13^. The lack of correlation between IRF7 and ethanol consumption in other regions involved in addiction behaviors, such as the PrL and Acb, suggest a region-specific role of IRF7 signaling in driving drinking behaviors. Similar to IRF7, we observed a region-specific trending increase in Iba-1 expression in the aIC following repeated CIE and ethanol self-administration, which was not due to an increase in microglial number. These data indicate aIC microglia maintain a pro-inflammatory state 72 hours into abstinence. To better understand if this microglial phenotype was contributing to CIE-induced IRF7 expression we exposed rats to one cycle of CIE with and without microglial inhibition via minocycline. Unlike our findings after repeated CIE, we did not find ethanol-related influence on Iba-1 expression after one cycle of CIE, suggesting repeated CIE may be necessary to observe morphological microglial changes in early abstinence. However, microglial inhibition via minocycline significantly decreased ethanol-related IRF7 levels in both the aIC and Acb after one cycle of CIE. These data suggest that microglia pro-inflammatory signaling has a role in orchestrating molecular phenotypes related to ethanol self-administration during early abstinence. Therefore, profiling of microglia beyond Iba-1 expression is necessary to understand their function in early abstinence, leading us to perform LC-MS/MS characterization of microglia.

LC-MS/MS analysis indicates a pro-inflammatory microglial state 72 hours after one cycle of CIE. Using IPA, we find CIE led to a predicted increase in NLR and IL signaling, class 1 MHC presentation, and endosomal TLR trafficking, phenotypes that have been associated with ethanol-related microglia activation in the past^27–29^. The strongest predicted pathways suppressed by CIE deal with suppressed mitochondrial respiration driven by reduced expression of electron transport chain proteins (i.e., ATP5B, COX6C). Previous studies find macrophages shift their metabolic state away from oxidative phosphorylation to promote inflammatory signing^24^, further suggesting a pro-inflammatory microglial phenotype due to CIE. Primary microglia from CIE-exposed rats also had predicted increase in translation pathways due to increased abundance of ribosomal proteins (RPS25) and regulators of the integrated stress response (i.e. EIF2S1, EIF3A), suggesting proteostasis stress^30^. Upregulation of translation pathways were also found in proteomics datasets from bulk OFC tissue from human male donors with AUD^23^, whereas metabolic pathways from human datasets were incongruent with the metabolic phenotype of rat aIC microglia following CIE. Further characterization of the microglia-specific proteome from human AUD donors is necessary to understand if the metabolic phenotype found here extends to humans.

Minocycline treatment at the time of CIE promoted oxidative phosphorylation and suppressed translation pathways. However, this signature was not observed due to minocycline alone, suggesting minocycline in combination with CIE opposes microglial stress pathways. Functionally it appears CIE may suppress the ability of microglia to remodel synapses, as indicated by down regulation of GO pathways dealing with synaptic vesicles, synaptic transmission, and transmembrane transportation. Synaptic pathways were predicted to be increased with the addition of minocycline, along with cellular movement pathways, suggesting a pro-surveillance, tissue remodeling microglial phenotype. Warden et al. 2020 also finds significant alterations in synaptic pathways in mPFC tissue from ethanol-dependent mice following microglia depletion^14^, further indicating microglial remodeling of neuronal synapses following ethanol exposure. Our data suggest microglial surveillance and modulation of aIC synapses may be necessary to suppress ethanol self-administration. Alternatively, recent data suggest microglia may provide endogenous neurotransmitters to neurons to modulate their function^31^. However, we did not detect changes to neuronal activation marker FosB due to minocycline treatment. Future studies may explore synaptic vesicle release from microglia or activation of specific neuronal subpopulations in the aIC following CIE and minocycline to better understand these protective mechanisms. Additionally, further work is needed to understand if the metabolic and ribosomal phenotypes indicated by IPA are related to predicted functional changes.

We find that minocycline treatment at the time of CIE reduced ethanol-self administration 72 hours into abstinence. These data suggest the aforementioned microglial phenotypes of early abstinence contribute to ethanol self-administration. Further, these data indicate that blocking the formation of these microglial phenotypes can suppress drinking during early abstinence. Following the self-administration task, rats were exposed to an additional cycle of CIE and sacrificed 72 hours later to profile microglial and IRF7 expression in early abstinence. Minocycline reduced microglial area in the aIC and Acb, both of which were positively correlated with increased ethanol consumption. Although we did not observe changes in aIC IRF7 expression due to minocycline, we replicated the finding that aIC IRF7 expression positively correlates with ethanol self-administration in aIC specific manner. We have reported previously that microglia secrete endogenous agonists of TLR7^19^, which is upstream of IRF7 activation. Since IRF7 protein is predominantly neuronal^13^, this suggests that microglial-neuronal communication contributes to escalation of drinking during abstinence.

In conclusion, these data establish a role for microglia in ethanol-self-administration escalation during early abstinence consistent with a cascade including microglia signaling and neuronal IRF7 induction leading to ethanol consumption. LC-MS/MS profiling of microglia during the early abstinence phase suggests rescuing CIE-related suppression of oxidative phosphorylation, induction of protein stress responses, or impaired functional regulation of synapses, may be viable targets to normalize microglial function and reduce ethanol-self administration. In this study, minocycline was administered during CIE and microglia were assessed during early abstinence. Although these data reveal microglial phenotypes present during the ethanol consumption phase, it is difficult to parse potential insufficient neuroprotective responses of microglia due to CIE versus determinantal responses. Future studies will investigate the ability to reverse dependence-induced increases in ethanol self-administration by targeting these processes in microglia.

### Funding

This work was funded by NIAAA AA011605 (LGC, JB) and NIAAA T32 AA007573 (PA)

## Methods

### Animals

Animals were kept under the care of the Division of Comparative Medicine at UNC-Chapel Hill and all procedures followed NIH Guide to Care and Use of laboratory Animals guidelines. Adult male Wistar rats (Charles Rivers Laboratories, Wilmington, MA, USA) were house in ventilated cages and given access to food and water ad libitum. Rats were kept on a 12hr light/dark cycle (7am/7pm) in a temperature and humidity-controlled room.

### Chronic-intermittent ethanol vapor (CIE) exposure

Rats underwent chronic intermittent ethanol vapor cycles (16hr/day 5:00 PM-9:00 AM) across four consecutive days (Monday PM to Friday AM). Two rats were placed in each ethanol exposure chamber, while air controls remained in home cages in the vivarium. 95% ethanol v/v was vaporized into fresh air and dispersed into each cage at a flow rate of ~16 L/min through an air compressor. Following each vapor session, blood was collected via tail snip and samples were centrifuged to obtain plasma. Plasma was analyzed using an AM1 Alcohol Analyzer (Analox Instruments Ltd, Stourbridge, UK). Adjustments were made to maintain blood ethanol levels in the 150-220mg/dL range.

### EtOH Self-administration

Ethanol self-administration was assessed in 30 min sessions in operant chambers (Med Associated Inc., St. Albans, VT) as previously described^13^. Each chamber was placed in a sound attenuating box with proper ventilation to mask outside noise. Each chamber had two retractable levers on the left and right walls. Levers were positioned under a cue light and beside a liquid delivery receptacle. Responses on the active lever (fixed-ratio 2; FR2 schedule) delivered 0.1 mL of liquid reinforcer; iAcbtive lever presses had no consequence. Rats were trained to self-administer ethanol via a sucrose-fading procedure, starting with 10% (w/v) sucrose + 2% (v/v) ethanol (10S/2E) during an overnight session. Daily self-administration sessions followed with the following progression of solutions: 10S/2E, 10S/5E, 10S/10E, 5S/10E, 5S/15E, 2S/15E, 2S/20E, 20E, and finally 15% ethanol (15E) which was maintained for the remainder of self-administration experiments.

### Perfusion and tissue collection

Rats were sacrificed and subjected to transcardial perfusion with 0.1 M phosphate-buffered saline (PBS, pH 7.4). Following perfusion, brains were excised and drop-fixed in 4.0% paraformaldehyde for immunohistochemical assessments. Coronal sections were cut (40 µm) on a sliding microtome (MICROM HM450; ThermoScientific, Austin, Texas, USA), and sections were sequentially collected into well plates and stored at −20°C in cryoprotectant (30% glycol/30% ethylene glycol in PBS).

### Immunofluorescence

Immunofluorescence was performed using our previously published protocols^11^. Briefly, free floating tissue sections were washed in 1x phosphate-buffered saline (PBS) and incubated in citrate buffer (Fisher Scientific, Waltham, MA; NC9935936). for antigen retrieval for 1 hour at 70 °C (Millipore Sigma, Burlington, MA; T8787). Sections were then blocked in a solution containing 4% normal goat serum and 0.1% Triton X-100. Sections were then incubated overnight in IRF7 (1:500, Biorbyt orb6232) or Iba-1 (Wako 019-19741) antibodies. The next day, sections were washed and incubated with fluor-conjugated secondary antibodies (1:1000, Invitrogen), washed, and mounted with Prolong Gold Anti-Fade mounting medium (Thermo Fisher Scientific; P36971). 3-4 sections were stained per animal. mages were taken on the Keyence BZ-X800 Microscope and analyzed using the accompanying BZ-X800 Analyzer software. Tissue sections were chosen based on desired bregma according to the rat brain atlas (Paxinos and Watson) (anterior insula [aIC, +3.00mm], nucleus accumbens core [Acb, +2.04mm], and the prelimbic cortex [PrL, +3.00mm]. Positive staining was measured as immunoreactive pixels/mm^2^.

### Primary Microglia Isolation

Microglia were isolated as previously described^32^. Rats were anesthetized and transcardially perfused with 1xPBS. Brains were removed and anterior insula punches were taken from both hemispheres. aIC tissue was then placed in an enzymatic digestion buffer (Hanks Balanced Salt Solution without magnesium or calcium [HBSS; ThermoFisher, #14175095], 5% fetal bovine serum [FBS; ThermoFisher, #A3160501], 10 µM HEPES [ThermoFisher, #15630080], 2.0 mg/mL collagenase A [Millipore, #10103586001], and 28 U/mL DNase I [Millipore, #10104159001]) at 37C for 45 minutes. Every 15 minutes brain tissue was passed through successively smaller pasture pipettes to generate single cell suspensions. Cell suspensions were passed through 70uM cell strainers and centrifuged (300xg,). Debris removal (Miltenyi #130-109-398) and Cd11b+ selection was performed to isolate primary microglia according to manufacturer’s instructions (Miltenyi #130-105-643). Microglia were then washed three times with 1x PBS, pelleted, snap frozen, and stored at −80C prior to LC-MS/MS.

### LC-MS/MS proteomic analysis of isolated microglia

Microglial LC-MS/MS was run as previously described^32^. Microglia (n=4/group) were reduced with 5mM DTT at 37ºC for 45 min then alkylated with 15mM iodoacetamide at RT for 45 min, shielded from light. Samples were then diluted to 1 M urea with 50 mM ammonium bicarbonate and subjected to digestion with trypsin (Promega) overnight at 37ºC at a 1:50 enzyme: protein ratio. Peptides were then acidified to 0.5% trifluoroacetic acid and desalted using ZipTips (Sigma) followed by vacuum centrifugation. Elutes were then resuspended in 2% acetonitrile with 0.1% formic acid. To perform LC-MS/MS the peptide samples were analyzed by Easy nLC 1200 coupled to a QExactive HF mass spectrometer (Thermo Scientific). Samples were injected onto an Aurora Ultimate TS column (75 μm id × 25 cm, 1.7 μm particle size) (IonOpticks) and separated over a 90-minute period. The gradient for separation consisted of 5–45% mobile phase B at a 250 nl/min flow rate, where mobile phase A was 0.1% formic acid in water and mobile phase B consisted of 0.1% formic acid in 80% ACN. The QExactive HF was operated in data-dependent mode where the 15 most intense precursors were selected for subsequent fragmentation. Resolution for the precursor scan (m/z 350–1700) was set to 60,000, while MS/MS scans resolution was set to 15,000. The normalized collision energy was set to 27% for HCD. Peptide match was set to preferred, and precursors with unknown charge or a charge state of 1 and ≥□7 were excluded.

For data analysis, raw data files were searched against the Uniprot reviewed and unreviewed rat database appended with a contaminants database, using the Sequest HT search engine node within Proteome Discoverer (v3.1, Thermo Fisher). Enzyme specificity was set to trypsin, up to two missed cleavage sites were allowed, methionine oxidation and N-terminus acetylation were set as variable modifications and cysteine carbamidomethylation was set as a static modification. The Minora node was used to extract label-free quantification (LFQ) intensities. A 1% peptide-level false discovery rate (FDR) and a 1% protein-level FDR was used to filter all data. Match between runs was enabled, and a minimum of two peptides was required for label-free quantitation using the LFQ intensities. Perseus was used for further processing [62]. Proteins with >□50% of missing values across the samples were filtered out. The remaining missing values were imputed from normal distribution within Perseus. Log2 fold change (FC) ratios were calculated using the averaged Log2 LFQ intensities of microglial repopulation or amyloid to control, and students t-test performed for each pairwise comparison, with p-values calculated. Proteins with p-values (<□0.05) and Log2FC□>□0.5 or Log2FC <-0.5 were considered significant.

### IPA and GO analysis

To identify the influence of CIE and/or minocycline on biological pathways, protein identifiers, log2fc values, and experimental p-values were processed by Qiagen Ingenuity Pathway Analysis (IPA). Through IPA’s canonical pathways analysis, biological pathways were predicted to be up or down regulated as signified by z-score and p-value. We also employed Gene Ontology (GO) analysis for further classification of protein functions given our treatment conditions. Protein identifiers, log2fc, and p-values were processed in R (version 4.4.1) using the cluster::profiler::gseGO package^33^.

### Statistical Analysis

Statistical analyses were performed on Graphpad prism (10.2.2). For preplanned comparisons between two groups, unpaired two-tailed Student *t-*tests were employed. A *p*□*<*□0.05 was considered as significant. For comparisons with three groups, 1-way ANOVAs were used with Sidak’s post-hoc testing when appropriate. Outliers were identified using Grubb’s test (GraphPad).

## Supplemental Figure legends

**Supplementary Figure 1.**
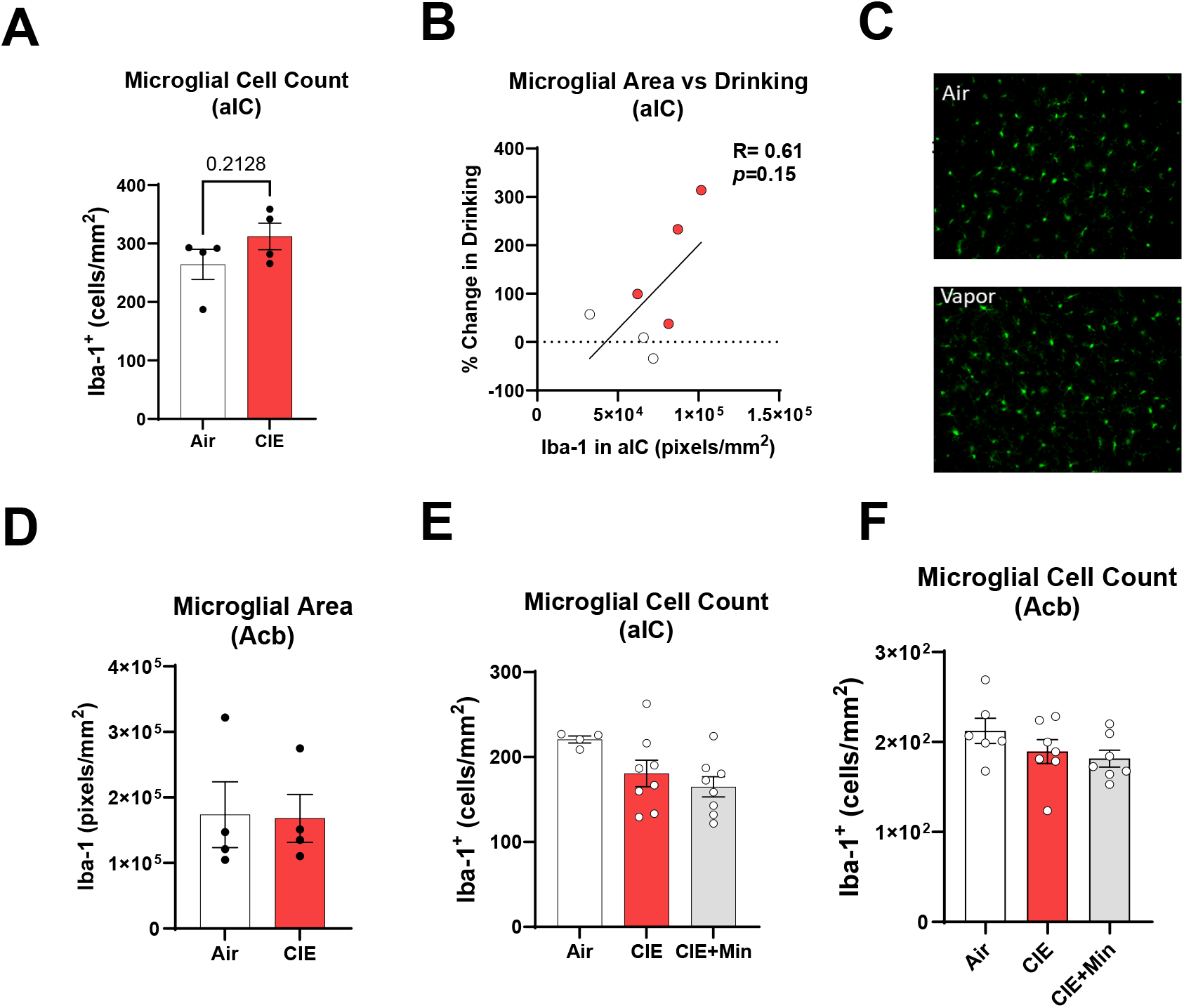
No differences in microglia numbers due to CIE or minocycline. (A) Microglia cell count in the aIC following multiple rounds of CIE and self-administration. (B) Correlation between ethanol self-administration escalation and aIC Iba-1 levels. Red dots indicate CIE, white dots indicate Air group. (C-D) Iba-1 expression in the Acb following CIE and ethanol self-administration. Microglia cell count in the (E) aIC and (F) Acb following one cycle of CIE and minocycline treatment.

**Supplementary Figure 2.**
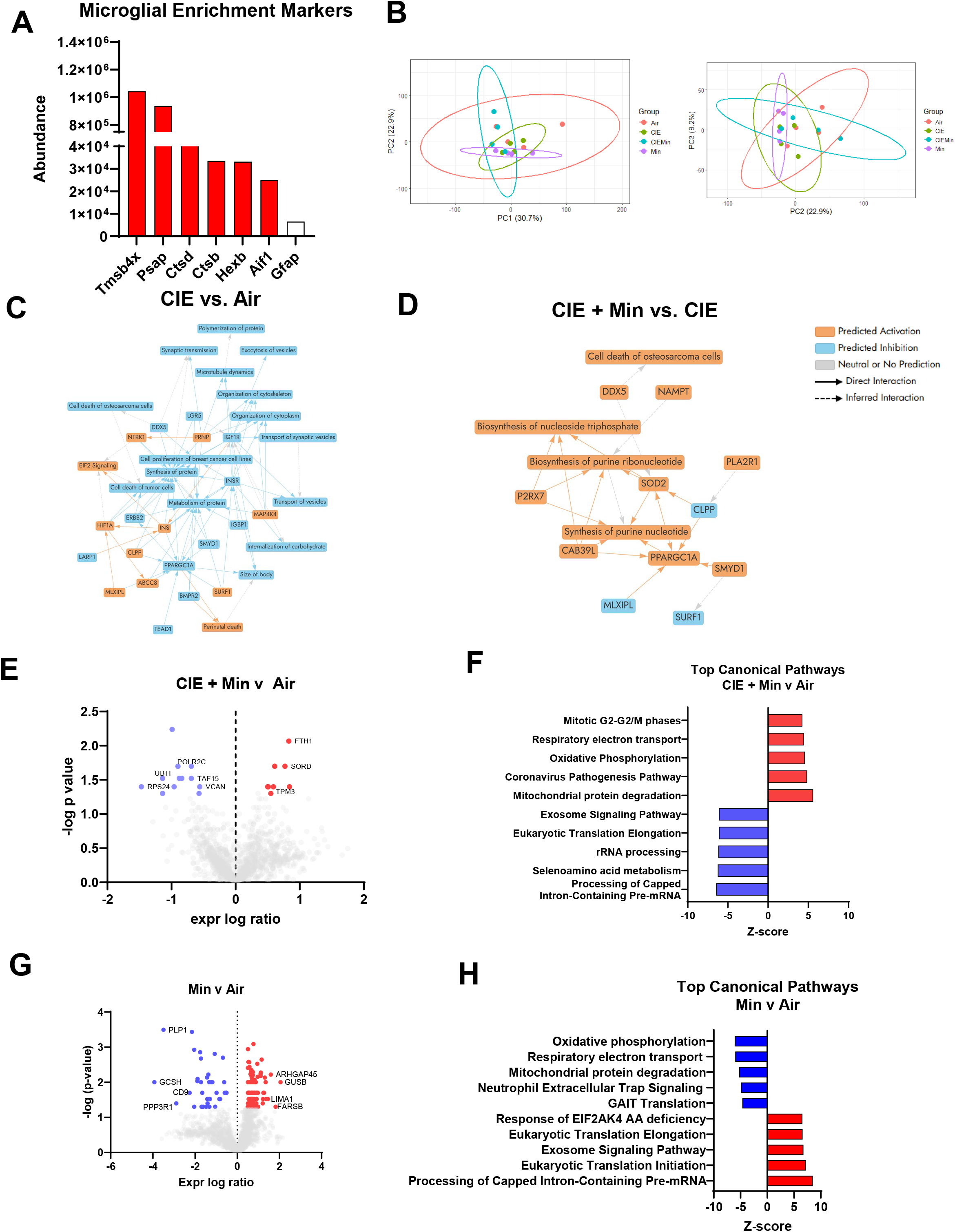
Isolation and LC-MS/MS profiling of aIC microglia. (A) Abundance of microglia specific proteins (red bars) and astrocyte specific proteins (white bar) following Cd11b+ selection. (B) PCA plots of aIC microglia proteome from Air, Min, CIE, CIE+Min groups. IPA network analysis of (C) CIE v Air comparison and (D) CIE + Min v CIE comparison. (E) Volcano plot of significantly differentially expressed proteins from the CIE + Min v Air comparison. (F) Top 5 up and down regulated IPA pathways from the CIE + Min v Air comparison. (G) Volcano plot of significantly differentially expressed proteins from the Min v Air comparison. (H) Top 5 up and down regulated IPA pathways from the Min v Air comparison.

**Supplementary Figure 3.**
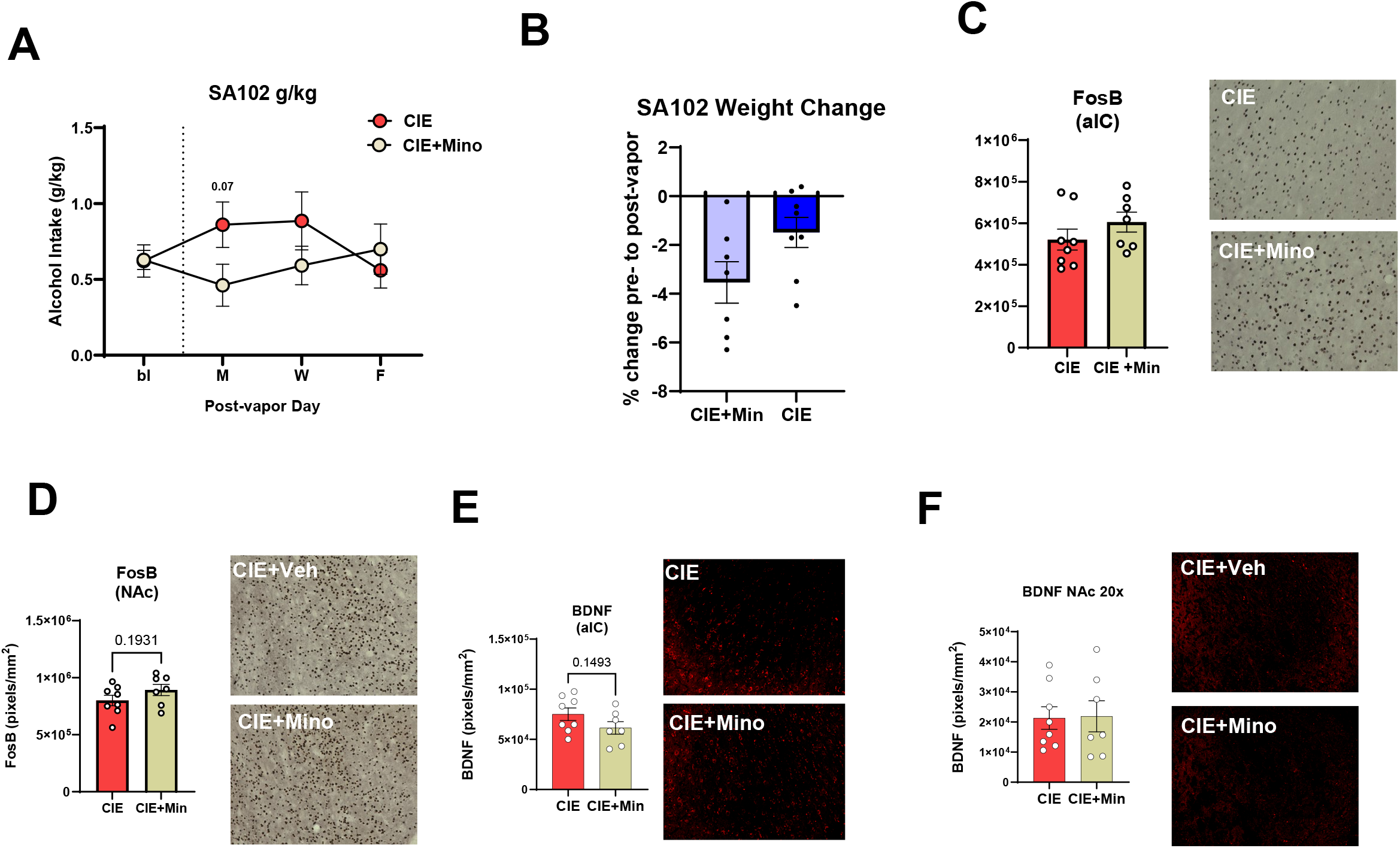
Minocycline does not influence BDNF or FosB in the aIC. (A) Ethanol intake (g/kg) during each self-administration day following one cycle of CIE and minocycline or vehicle treatment. (B) No change in weight observed due to minocycline treatment. FosB expression in the (C) aIC and (D) Acb following CIE and minocycline. Immunofluorescence for total and microglial BDNF expression in the (E) aIC and (F) Acb. N=4/group.

## References

1. Alcoholism., N.I.o.A.A.a. (August 15, 2025.). Alcohol Facts and Statistics.

2. Carr, T., Kilian, C., Llamosas-Falcón, L., Zhu, Y., Lasserre, A.M., Puka, K., and Probst, C. (2024). The risk relationships between alcohol consumption, alcohol use disorder and alcohol use disorder mortality: A systematic review and meta-analysis. Addiction 119, 1174–1187. 10.1111/add.16456.

3. Koob, G.F., and Volkow, N.D. (2010). Neurocircuitry of Addiction. Neuropsychopharmacology 35, 217–238. 10.1038/npp.2009.110.

4. Sinha, R. (2011). New findings on biological factors predicting addiction relapse vulnerability. Curr Psychiatry Rep 13, 398–405. 10.1007/s11920-011-0224-0.

5. Coleman, L.G., Zou, J., and Crews, F.T. (2020). Microglial depletion and repopulation in brain slice culture normalizes sensitized proinflammatory signaling. Journal of Neuroinflammation 17, 27. 10.1186/s12974-019-1678-y.

6. Crews, F.T., Lawrimore, C.J., Walter, T.J., and Coleman, L.G., Jr. (2017). The role of neuroimmune signaling in alcoholism. Neuropharmacology 122, 56–73. 10.1016/j.neuropharm.2017.01.031.

7. Pascual, M., Baliño, P., Alfonso-Loeches, S., Aragón, C.M.G., and Guerri, C. (2011). Impact of TLR4 on behavioral and cognitive dysfunctions associated with alcohol-induced neuroinflammatory damage. Brain, Behavior, and Immunity 25, S80–S91. 10.1016/j.bbi.2011.02.012.

8. Crews, F.T., Qin, L., Coleman, L., Vidrascu, E., and Vetreno, R. (2026). Cortical reactive microglia activate astrocytes, increasing neurodegeneration in human alcohol use disorder. Brain, Behavior, and Immunity 131, 106156. 10.1016/j.bbi.2025.106156.

9. Crews, F.T., Qin, L., Sheedy, D., Vetreno, R.P., and Zou, J. (2013). High mobility group box 1/Toll-like receptor danger signaling increases brain neuroimmune activation in alcohol dependence. Biological psychiatry 73, 602–612. 10.1016/j.biopsych.2012.09.030.

10. Qin, L., Zou, J., Barnett, A., Vetreno, R.P., Crews, F.T., and Coleman Jr, L.G. (2021). TRAIL mediates neuronal death in AUD: a link between neuroinflammation and neurodegeneration. International journal of molecular sciences 22, 2547.

11. McNair, E.M., Dawkins, L.W., Materia, B., Ross, G., Barnett, A., Nakkala, P., Qin, L., Zou, J., Nikolova, V., Moy, S., and Coleman, L.G. (2025). Microglia promote neurodegeneration and hyperkatifeia during withdrawal and abstinence from binge alcohol. The American Journal of Pathology. 10.1016/j.ajpath.2025.10.005.

12. Qin, L., He, J., Hanes, R.N., Pluzarev, O., Hong, J.-S., and Crews, F.T. (2008). Increased systemic and brain cytokine production and neuroinflammation by endotoxin following ethanol treatment. Journal of neuroinflammation 5, 10.

13. Lovelock, D.F., Carew, J.M., McNair, E.M., Materia, B.M., Darawsheh, S.Z., Downs, A.M., Sizer, S.E., McDonald, S.A., McElligott, Z.A., Coleman, L.G., Jr., and Besheer, J. (2026). Interferon-Regulatory Factor 7: A Neuroimmune Role for Vapor-Induced Escalations in Ethanol Self-Administration. bioRxiv. 10.64898/2026.04.01.715945.

14. Warden, A.S., Wolfe, S.A., Khom, S., Varodayan, F.P., Patel, R.R., Steinman, M.Q., Bajo, M., Montgomery, S.E., Vlkolinsky, R., Nadav, T., et al. (2020). Microglia Control Escalation of Drinking in Alcohol-Dependent Mice: Genomic and Synaptic Drivers. Biological Psychiatry 88, 910–921. 10.1016/j.biopsych.2020.05.011.

15. Kobayashi, K., Imagama, S., Ohgomori, T., Hirano, K., Uchimura, K., Sakamoto, K., Hirakawa, A., Takeuchi, H., Suzumura, A., and Ishiguro, N. (2013). Minocycline selectively inhibits M1 polarization of microglia. Cell death & disease 4, e525–e525.

16. Agrawal, R.G., Hewetson, A., George, C.M., Syapin, P.J., and Bergeson, S.E. (2011). Minocycline reduces ethanol drinking. Brain, Behavior, and Immunity 25, S165–S169. 10.1016/j.bbi.2011.03.002.

17. Lovelock, D.F., Liu, W., Langston, S.E., Liu, J., Van Voorhies, K., Giffin, K.A., Vetreno, R.P., Crews, F.T., and Besheer, J. (2022). The Toll-like receptor 7 agonist imiquimod increases ethanol self-administration and induces expression of Toll-like receptor related genes. Addiction Biology 27, e13176.

18. Qin, L., Zou, J., Barnett, A., Vetreno, R.P., Crews, F.T., and Coleman, L.G. (2021). TRAIL Mediates Neuronal Death in AUD: A Link between Neuroinflammation and Neurodegeneration. International journal of molecular sciences 22. 10.3390/ijms22052547.

19. Coleman, L.G., Jr., Zou, J., and Crews, F.T. (2017). Microglial-derived miRNA let-7 and HMGB1 contribute to ethanol-induced neurotoxicity via TLR7. Journal of neuroinflammation 14, 22. 10.1186/s12974-017-0799-4.

20. Vendruscolo, L.F., and Roberts, A.J. (2014). Operant alcohol self-administration in dependent rats: focus on the vapor model. Alcohol 48, 277–286. 10.1016/j.alcohol.2013.08.006.

21. O’Dell, L.E., Roberts, A.J., Smith, R.T., and Koob, G.F. (2004). Enhanced Alcohol Self-Administration after Intermittent Versus Continuous Alcohol Vapor Exposure. Alcoholism: Clinical and Experimental Research 28, 1676–1682. 10.1097/01.ALC.0000145781.11923.4E.

22. Crews, F.T., Qin, L., Coleman, L., Vidrascu, E., and Vetreno, R. (2025). Cortical reactive microglia activate astrocytes, increasing neurodegeneration in human alcohol use disorder. Brain Behav Immun 131, 106156. 10.1016/j.bbi.2025.106156.

23. Mulholland, P.J., Berto, S., Wilmarth, P.A., McMahan, C., Ball, L.E., and Woodward, J.J. (2023). Adaptor protein complex 2 in the orbitofrontal cortex predicts alcohol use disorder. Mol Psychiatry 28, 4766–4776. 10.1038/s41380-023-02236-3.

24. Devanney, N.A., Stewart, A.N., and Gensel, J.C. (2020). Microglia and macrophage metabolism in CNS injury and disease: The role of immunometabolism in neurodegeneration and neurotrauma. Exp Neurol 329, 113310. 10.1016/j.expneurol.2020.113310.

25. Fernández, D., Geisse, A., Bernales, J.I., Lira, A., and Osorio, F. (2021). The Unfolded Protein Response in Immune Cells as an Emerging Regulator of Neuroinflammation. Front Aging Neurosci 13, 682633. 10.3389/fnagi.2021.682633.

26. Witkiewitz, K., and Villarroel, N.A. (2009). Dynamic association between negative affect and alcohol lapses following alcohol treatment. J Consult Clin Psychol 77, 633–644. 10.1037/a0015647.

27. Alfonso-Loeches, S., Urena-Peralta, J., Morillo-Bargues, M.J., Gómez-Pinedo, U., and Guerri, C. (2016). Ethanol-induced TLR4/NLRP3 neuroinflammatory response in microglial cells promotes leukocyte infiltration across the BBB. Neurochemical research 41, 193–209.

28. Anton, P.E., Rutt, L.N., Kaufman, M.L., Busquet, N., Kovacs, E.J., and McCullough, R.L. (2024). Binge ethanol exposure in advanced age elevates neuroinflammation and early indicators of neurodegeneration and cognitive impairment in female mice. Brain, behavior, and immunity 116, 303–316.

29. Coleman Jr, L.G., Zou, J., and Crews, F.T. (2017). Microglial-derived miRNA let-7 and HMGB1 contribute to ethanol-induced neurotoxicity via TLR7. Journal of neuroinflammation 14, 22.

30. Hetz, C. (2012). The unfolded protein response: controlling cell fate decisions under ER stress and beyond. Nature Reviews Molecular Cell Biology 13, 89–102. 10.1038/nrm3270.

31. Sogn, C.J., Valori, M., Rønning, P., Eide, P.K., Gundersen, V., and Nordengen, K. (2026). Context-dependent localization and expression of glutamate and GABA and their machinery in microglia. Journal of Neuroinflammation.

32. Zou, J., McNair, E., DeCastro, S., Lyons, S.P., Mordant, A., Herring, L.E., Vetreno, R.P., and Coleman Jr, L.G. (2024). Microglia either promote or restrain TRAIL-mediated excitotoxicity caused by Aβ1− 42 oligomers. Journal of neuroinflammation 21, 215.

33. Korotkevich, G., Sukhov, V., Budin, N., Shpak, B., Artyomov, M.N., and Sergushichev, A. (2016). Fast gene set enrichment analysis. biorxiv, 060012.

